# Kinetic pathways of water exchange in the first hydration shell of magnesium: Influence of water model and ionic force field

**DOI:** 10.1101/2021.06.21.449344

**Authors:** Sebastian Falkner, Nadine Schwierz

## Abstract

Water exchange between the first and second hydration shell is essential for the role of Mg^2+^ in biochemical processes. In order to provide microscopic insights into the exchange mechanism, we resolve the exchange pathways by all-atom molecular dynamics simulations and transition path sampling. Since the exchange kinetics relies on the choice of the water model and the ionic force field, we systematically investigate the influence of seven different polarizable and non-polarizable water and three different Mg^2+^ models. In all cases, water exchange can occur either via an indirect or direct mechanism (exchanging molecules occupy different/same position on water octahedron). In addition, the results reveal a crossover from an interchange dissociative (I_*d*_) to an associative (I_*a*_) reaction mechanism dependent on the range of the Mg^2+^-water interaction potential of the respective force field. Standard non-polarizable force fields follow the I_*d*_ mechanism in agreement with experimental results. By contrast, polarizable and long-ranged non-polarizable force fields follow the I_*a*_ mechanism. Our results provide a comprehensive view on the influence of the water model and ionic force field on the exchange dynamics and the foundation to assess the choice of the force field in biomolecular simulations.

## Introduction

In aqueous solutions, Mg^2+^ ions are surrounded by a hydration shell of six water molecules, which is subsequently enclosed by a second hydration shell. Water exchange between these hydration shells plays an important role in a large variety of biochemical processes ranging from simple ion pair formation to catalyzed reactions in metalloenzymes or the transport of ions across cell membranes [1–5]. Since water exchange governs any type of reaction involving the replacement of the strongly bound hydration waters, resolving the reaction mechanism has received considerable scientific attention in experiments and simulations [6–13].

Experimental techniques such as dielectric relaxation, X-ray adsorption, femtosecond mid-infrared and far-infrared adsorption spectroscopy provide insight into the solvation structure of Mg^2+^ [2, 14–16] while NMR experiments facilitate the direct measurement of water exchange rates [6–8]. However, the structural changes during water exchange are experimentally not directly accessible. Still, according to the mechanistic classification for ligand exchange reactions proposed by Langford and Gray [17], the mechanism can be divided into four categories: associative (A), dissociative (D), interchange associative (I_*d*_) and interchange dissociative (I_*a*_). In the former two categories, a detectable intermediate with increased (A) or decreased (D) coordination number exists. For the latter two categories, no kinetically detectable intermediate exists. In order to discriminate between the four categories, the activation volume is typically used for the experimental identification of the water exchange mechanism [7, 18]. For Mg^2+^, the activation volume is positive and based on the similarity with Co^2+^ and Ni^2+^, the I_*d*_ or D mechanism has been proposed [7, 8]. However, the question, which mechanism is dominant could not be settled with certainty [8].

Here, simulations could provide further insights by a unique atomistic description of the exchange dynamics. However, simulating water exchange is tremendously challenging for two reason. (i) Water exchange around Mg^2+^ is rare. According to experiments water exchange is on the microsecond timescale [6–8]. Therefore, straight forward simulations are unsuitable to sample water exchange with sufficient statistics. (ii) Water exchange involves the concerted motion of several water molecules. The water molecules exchange in a concerted fashion and molecules in the first hydration shell and beyond rearrange collectively [12]. Consequently, ab initio quantum mechanical calculations are not feasible to resolve the exchange dynamics due to their high computational costs and limited system size. On the other hand, classical all-atom simulations in combination with transition path sampling have proven to be a particularly powerful sampling strategy to resolve the molecular exchange pathways [12]. However, the simulations rely on the accuracy of the available water and Mg^2+^ force fields. In particular, different water models and ionic force fields can have significant effects on the structural, thermodynamic and kinetic properties [10, 13, 19, 20]. For example, the rate of water exchange in the first hydration shell of Mg^2+^ varies by more than eight orders of magnitude depending on the exact choice of the force field [10, 13]. The choice of the water model and the ionic force field is therefore crucial to yield a quantitative description of Mg^2+^ in biomolecular simulations. In particular, capturing the mechanism of ligand exchange is essential to describe the role of Mg^2+^ in biochemical processes.

In order to assess the choice of different force fields in biomolecular simulations involving Mg^2+^, we here systematically investigate the influence of seven different polarizable and non-polarizable water and three different Mg^2+^ models. Using all-atom molecular dynamics simulations in combination with transition path sampling, we cover the long timescales involved and resolve the kinetic pathways. The results provide comprehensive insights into the influence of the water model and the ionic force field on the exchange mechanism.

## METHODS

### Atomistic model and simulation setup

The systems consist of one Mg^2+^ ion and 506 water molecules in a cubic simulation box (L=25 Å). We used three different polarizable and non-polarizable Mg^2+^ force fields and seven different water models as described below.

For the non-polarizable systems, the simulations were performed using Gromacs 2018.8 [21]. Long-range electrostatic interactions were accounted for using particle-mesh Ewald summation with a Fourier spacing of 0.12 nm and a grid interpolation up to order 4. Shortrange Coulomb and Lennard-Jones (LJ) interactions were cut off at 1.2 nm. Long-range dispersion corrections for energy and pressure were applied to correct for the truncated LJ potential. Prior to transition path sampling, the systems were minimized using the steepest descent algorithm. An NVT and NPT equilibration was performed each for 1 ns. All path sampling simulations were performed in the NVT ensemble at a temperature of 300 K with a time step of 2 fs using the velocity rescaling thermostat with stochastic term [22].

For the polarizable systems, the simulations were performed in OpenMM 7.4.1 [23]. Shortrange Coulomb and Lennard-Jones (LJ) interactions were cut off at 1.2 nm. Long-range electrostatic interactions were treated using particle-mesh Ewald summation. Simulations were performed with mutual polarization using a convergence criteria of 10^−5^ D. Trial trajectories were generated using the velocity Verlet with velocity randomization integrator [24] from the OpenMMTools library [25] at 300 K with a time step of 0.5 fs.

### Mg^2+^ force fields

We used three different Mg^2+^ force fields: The non-polarizable Mg^2+^ by Mamatkulov et al. [20], the microMg parameters [13], and polarizable Mg^2+^ parameters [26, 27] from the AMOEBA-2013 force field [28, 29]. The Mamatkulov Mg^2+^ parameters, were optimized in our previous work [20] to reproduce the experimental solvation free energy and the activity derivative. To test the influence of the water, we here used the Mamatkulov Mg^2+^ parameters in combination with six different non-polarizable water models, described in the corresponding section of the manuscript. In addition, we use the microMg parameters in combination with the TIP3P water model. The microMg parameters were optimized in our most recent work [13] to reproduce the experimental water exchange rate in addition to the before-mentioned thermodynamic properties. Finally, we used the polarizable AMOEBA force field [28, 29] with optimized Mg^2+^ parameters [26, 27] to explicitly account for the polarizability of Mg^2+^ and water.

### Water models

We used seven different non-polarizable and polarizable water models. From the different three site models, we chose the two most commonly used models, namely the SPC/E [30] and the TIP3P [31] water model. From the different four site models, we chose TIP4P/2005 [32] and TIP4P-D [33]. TIP4P/2005 has gained popularity and is often quoted as the best nonpolarizable general-purpose model [34]. TIP4P-D is one of the newer offsprings in the TIP4P family and was designed to improve water dispersion interactions. From the different five site models, we chose TIP5P-E [35] and TIP5P/2018 [36]. TIP5P-E is the re-parameterized version of the original TIP5P model [37] for use with Ewald summation methods. Since the TIP5P water model has not performed up to the initial expectations [34], we also used the TIP5P/2018 water model due to its improved bulk properties and its expected good performance in biomolecular simulations. As polarizable water model, the AMOEBA water model [28, 29] was used.

### Transition path sampling and transition state ensemble

Transition path sampling in the non-polarizable system was performed using OpenPathSampling [38, 39] with Gromacs 2018.8 [21]. An initial path was created using constant force pulling along the Mg^2+^-*O*_*w*_ distance using PLUMED [40]. New trial trajectories were created by selection of a snapshot, velocity randomization and integration forward and backward in time. For path sampling with Mg^2+^ parameters from Mamatkulov et al. [20], the shooting point selection was biased along the Mg^2+^-*O*_*w*_ distance with respect to the leaving water molecule using a Gaussian centered at 0.325 nm with and a width of 150 nm^−2^ [41]. The path length was flexible and the integration stopped when a stable state was reached. The states were defined based on the Mg^2+^-*O*_*w*_ distance with respect to the leaving water molecule. Additionally, the number of water oxygen atoms in the inner-shell with a cutoff of 0.21 nm was included to ensure that Mg^2+^ is coordinated by six water molecules in the stable states.

In the polarizable system, transition path sampling was performed using OpenPathSampling [38, 39] with OpenMM 7.4.1 [23]. Here, the initial path was generated at high temperature. As in the non-polarizable setup, new trial trajectories were generated by two-way shooting with randomized velocities. In contrast, it was not necessary to impose a bias on the snapshot selection. As previously, the stable states were defined based on the Mg^2+^-*O*_*w*_ distance with respect to the leaving water molecule. Due to the different exchange mechanism (I_*a*_), the number of water oxygen atoms within a cutoff of 0.35 nm was used to ensure Mg^2+^ is coordinated by six water molecules when the stable states are reached. The same state definitions were used in combination with the non-polarizable simulation protocol for the sampling of exchanges with the microMg parameters. For both non-polarizable and polarizable setups, the sampling was performed until 2500 trials were accepted.

Based on our previous work [12], we identify transition states as configurations that fulfill the criterion |*r*_1_ *− r*_2_| *<* 0.025 nm. Here, the incoming and leaving water molecules are approximately equally distant from the Mg^2+^ ion and the commitment probability to either stable state is similar. While efficient to evaluate, the criterion reproduces TIP3P transition state properties from our previous work [12].

### Active volume calculation

The activation volume was estimated as proposed by Qian et al. [42]. Van-der-Waals radii of atoms involved in the exchange process were scaled by 1.186. The solvent-excluded surface formed by these atoms was modeled using NanoShaper [43]. The volume enclosed by the solvent-excluded surface was estimated using Trimesh 3.8.8. A reference volume was obtained based on a 10 ns MD simulation without any exchange events. The activation volume defined as the change of volume with respect to the reference was then calculated for all transition states.

## RESULTS AND DISCUSSION

The aim of this work is to resolve how different water models and ionic force fields influence the kinetic pathways of water exchange in the first hydration shell of Mg^2+^. In the exchange reaction, one of the six water molecules from the first hydration shell is replaced by a water from the second one. To gain clear mechanistic insight, transition path sampling [44, 45] is applied to obtain a large number of independent reactive pathways. The advantage of transition path sampling is that it covers the up to millisecond long timescales involved [6, 8, 12] while generating an ensemble of true dynamic trajectories free of any bias [46]. This in turn allows us to unambiguously determine the reaction mechanism with sufficient statistics.

Four representative pathways are shown in Figure 1 corresponding to an associative or dissociative exchange pathway. Based on the distances between Mg^2+^ and the two exchanging water molecules, *r*_1_ and *r*_2_, associative and dissociative pathways are clearly distinguishable (Figure 1B). In the following, we show that the exchange occurs outside of the first hydration shell in dissociative pathways. During activation, one water molecule from the second hydration shell enters the molecular void leading to the concerted motion of another water molecule out of the first hydration shell. The distances of the entering and leaving water in the transition state are elongated and the activation volume is positive. By contrast, in associative pathways, the exchange occurs inside the first hydration shell, the distances are shortened and the activation volume is negative.

**FIG. 1:**
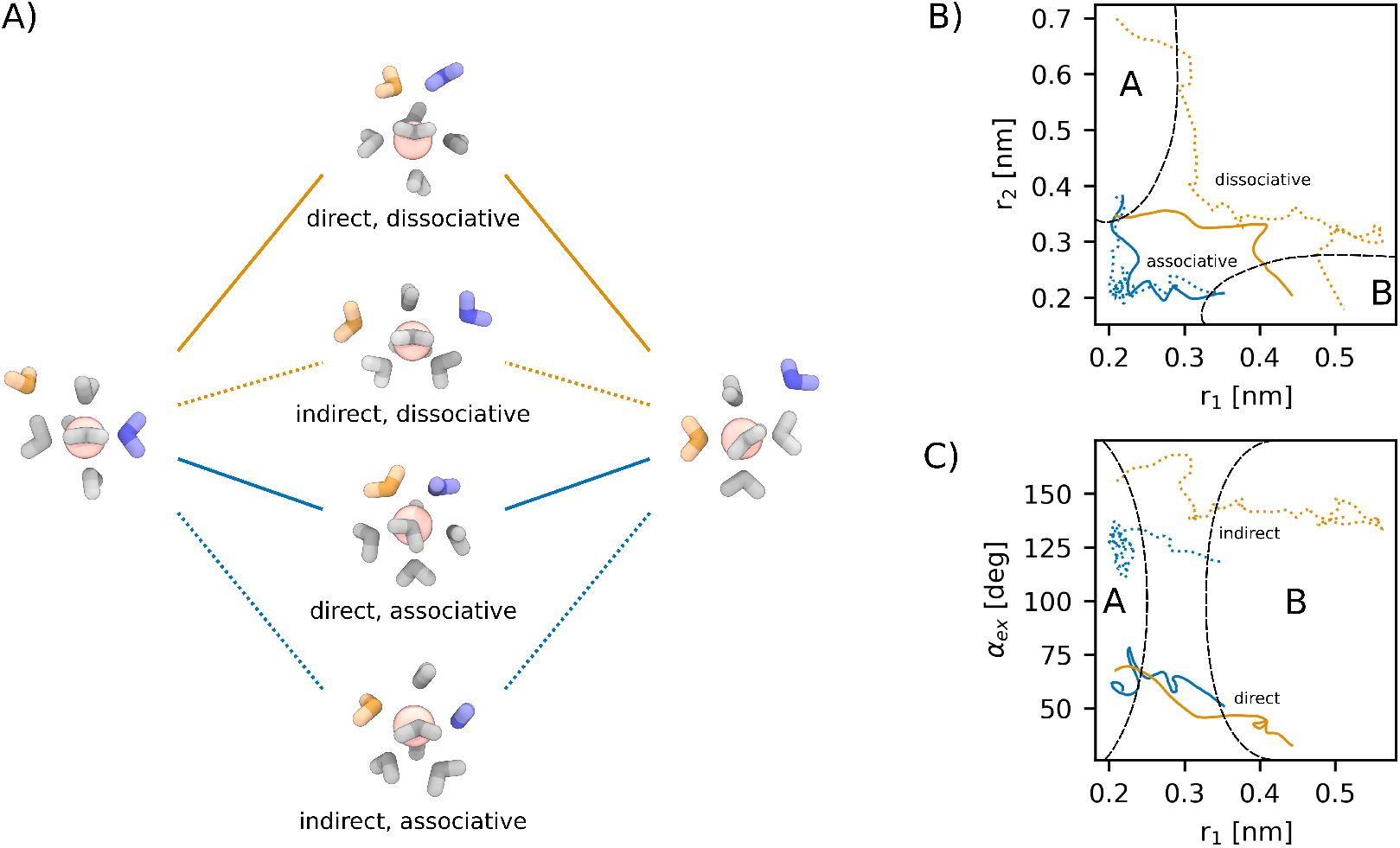
Overview of the observed exchange mechanisms for water exchange in the first hydration shell of Mg^2+^. (A) Simulations snapshots of the two stable states and the transition states for the spectrum of water exchange processes. (B) Four representative pathways connecting the two stable states as function the distance of the exchanging water molecules, *r*_1_ and *r*_2_. In this representation, associative and dissociative pathways are clearly separated. (C) Pathways as function of *r*_1_ and the exchange angle *α*_ex_ showing a clear separation of direct and indirect exchange mechanism.

As shown in Figure 1C, the exchange can be further classified by the exchange angle between incoming and outgoing water [12]. In the indirect exchange mechanism, entering and leaving water molecules occupy different positions on the water octahedron. In the direct exchange mechanism, the exchange takes place via an attack onto the edge of the water octahedron and the exchanging waters occupy the same positions on the water octahedron (Figure 1A). For instance in TIP3P, exchange angles *α*_ex_ *<* 90 correspond to the direct exchange mechanism while *α*_ex_ *>* 90 correspond to the indirect exchange mechanism (Figures 2B,C).

**FIG. 2:**
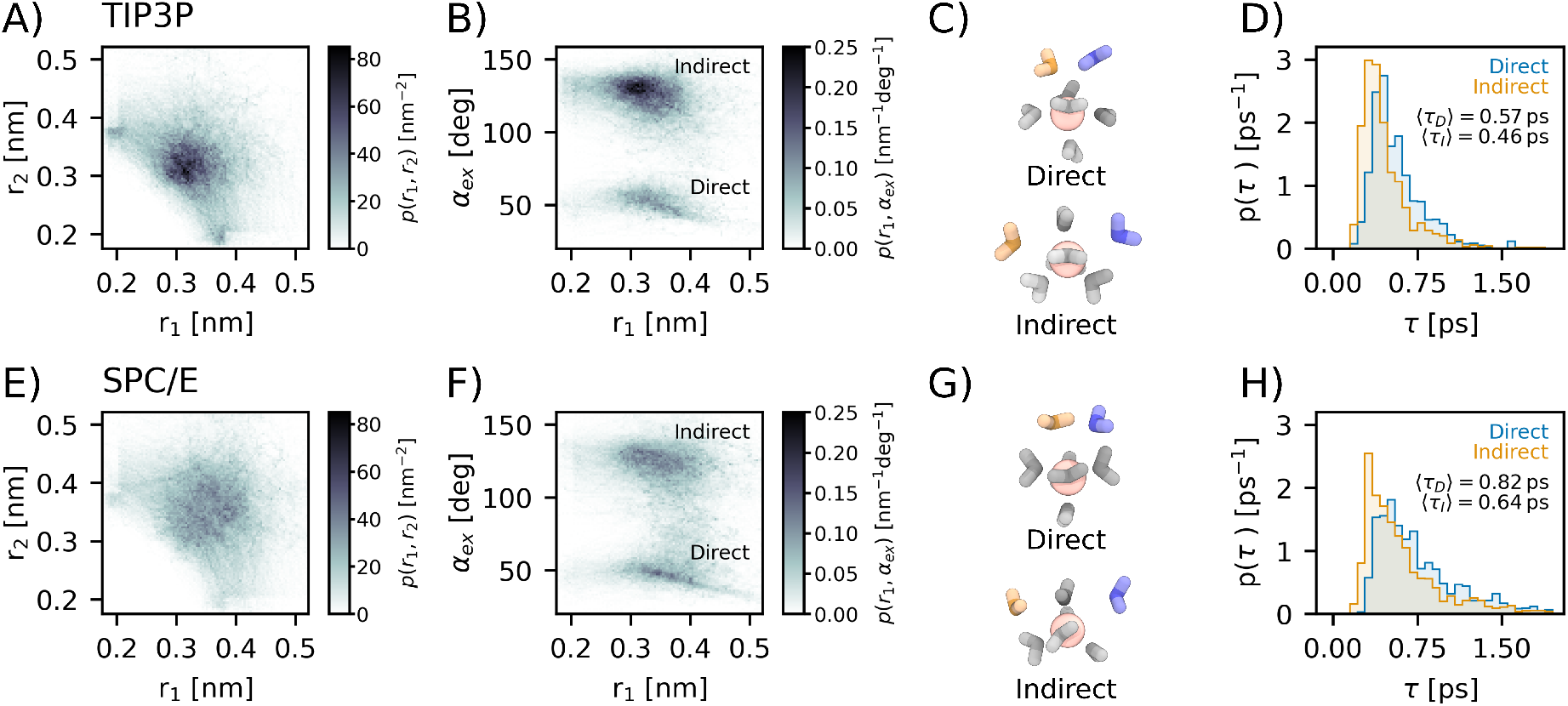
Water exchange with the three-site water models TIP3P (top) and SPC/E (bottom). (A,E) Probability distribution function *p*(*r*_1_, *r*_2_) along transition pathways in dependence of the distances *r*_1_ and *r*_2_. (B, F) Probability distribution function *p*(*r*_1_, *α*_ex_) in dependence of *r*_1_ and the exchange angle *α*_ex_. (C,G) Representative simulation snapshots of the transition state for the direct and indirect mechanism. (D, H) Distribution of transition times *p*(*τ*) for direct and indirect exchange pathways.

In the following, we provide insight into the exchange pathways obtained for seven water models and three different Mg2+ force fields. The probability distributions of the distances and exchange angle along reactive pathways provide direct insight into the kinetic pathways and the regions of high probability density coincide with the transition states (Figures 2-5, Table I).

**TABLE I:**
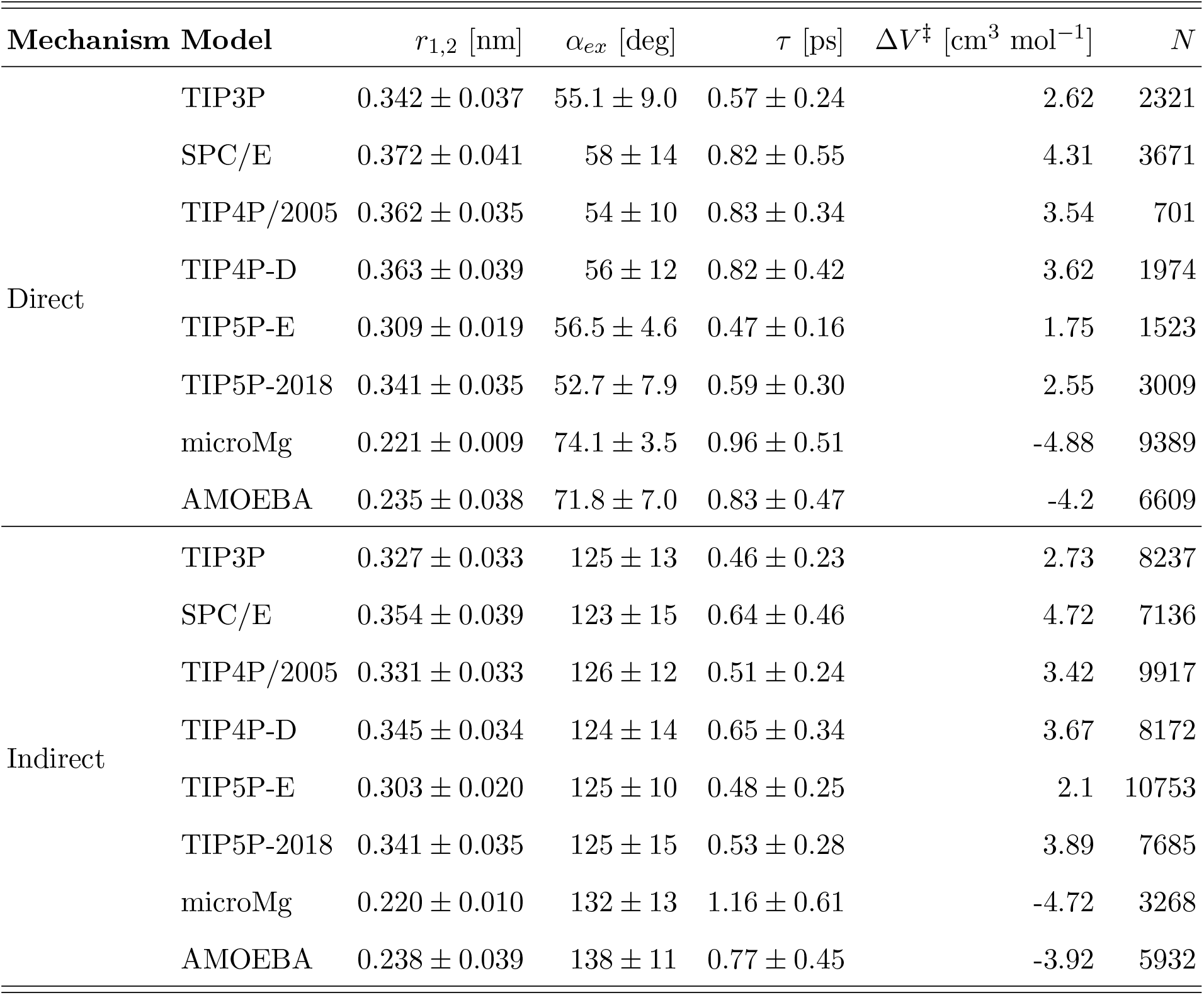
Properties of the transition state ensemble for direct and indirect exchange for different water models: Mg^2+^-oxygen distance of the exchanging water molecules, *r*_1,2_, angle between the exchanging water oxygens and Mg^2+^, *α*_*ex*_, average path length *τ*, activation volume Δ*V* ^‡^ and number *N* of direct and indirect exchange pathways. Averages and standard deviations of the properties are shown.

### Interchange dissociative pathways for rigid, non-polarizable, three, four and fivesite water models

Figure 2A, B shows the distributions of distances and exchange angles along reactive pathways in TIP3P water. In the transition state, the distances of the leaving and entering water molecules are elongated (Table I) and the exchange takes place outside the first hydration shell. In agreement with our previous results [12], two alternative exchange pathways, corresponding to the indirect and direct mechanism, can be clearly identified from the exchange angle (Figure 2B). In the direct exchange, the five spectator water molecules in the first hydration shell adopt a square pyramidal geometry in the transition state (Figure 2C). The indirect exchange occurs via a trigonal bipyramidal transition state. The exchanging water molecules enter and leave on opposing sides of the trigonal base (Figure 2C). This results in an exchange angle of approximately 120° (Table I). The activation volume is positive and larger *r*_1,2_ distances leads to a higher activation volume. Yet, this is not the only contributing factor. While indirect exchange reactions typically occur at smaller *r*_1,2_ distances than direct exchange reactions, they tend to have larger activation volumes. This is likely caused by the different angular distribution of the exchanging water in trans or cis position (Figure 2C). The distribution of transition times (Figure 2D) reveals that the water molecules spend less than 1 ps in transition and the indirect exchange is on average slightly faster compared to the direct exchange.

In summary, the leaving water molecule departs stepwise while an incoming water is approaching. The exchanging waters move in a concerted fashion with no kinetically detectable intermediate characteristic of an interchange (I) mechanism. Since the intrinsic activation volume is positive, direct and indirect water exchange corresponds to an interchange dissociative (I_*d*_) process in agreement with experimental results [7, 8]. It should be noted, however, that in equilibrium the indirect exchange is expected to be observed much more frequently compared to the direct mechanism since conformations with cis positions of exchanging ligands (direct pathways) are energetically less favorable compared to trans positions (indirect pathway) [9].

For SPC/E, TIP4P/2005, TIP4P-D, and TIP5P/20082018 the characteristics of water exchange remain essentially unchanged (Figures 2-4). However, for SPC/E the distribution of exchange distances (Figure 2E) is broader indicating that solvent reorientation plays a more pronounced role. This likely reflects the ideal tetrahedral shape of the SPC/E water model. Moreover, for TIP4P/2005, the indirect pathway almost completely disappears indicating that the cis position of the two exchanging water becomes less favorable (Figure 3B). The most pronounced differences are observed for TIP5P-E (Figure 4A,B). Here, the exchange takes place on the edge of the first hydration shell. The exchange distances are significantly shorter and the activation volume smaller compared to the other water models while the exchange angles remain largely unaffected (Table I). Consequently, I_*d*_ and I_*a*_ exchange mechanisms are almost indistinguishable since bond formation and bond breaking become equally important. Here, the different angular distribution of TIP5P-E may explain the observed differences [47]. In particular the marked difference to act as tetrahedral hydrogen bond donor and acceptor likely leads to an interchange mechanism.

**FIG. 3:**
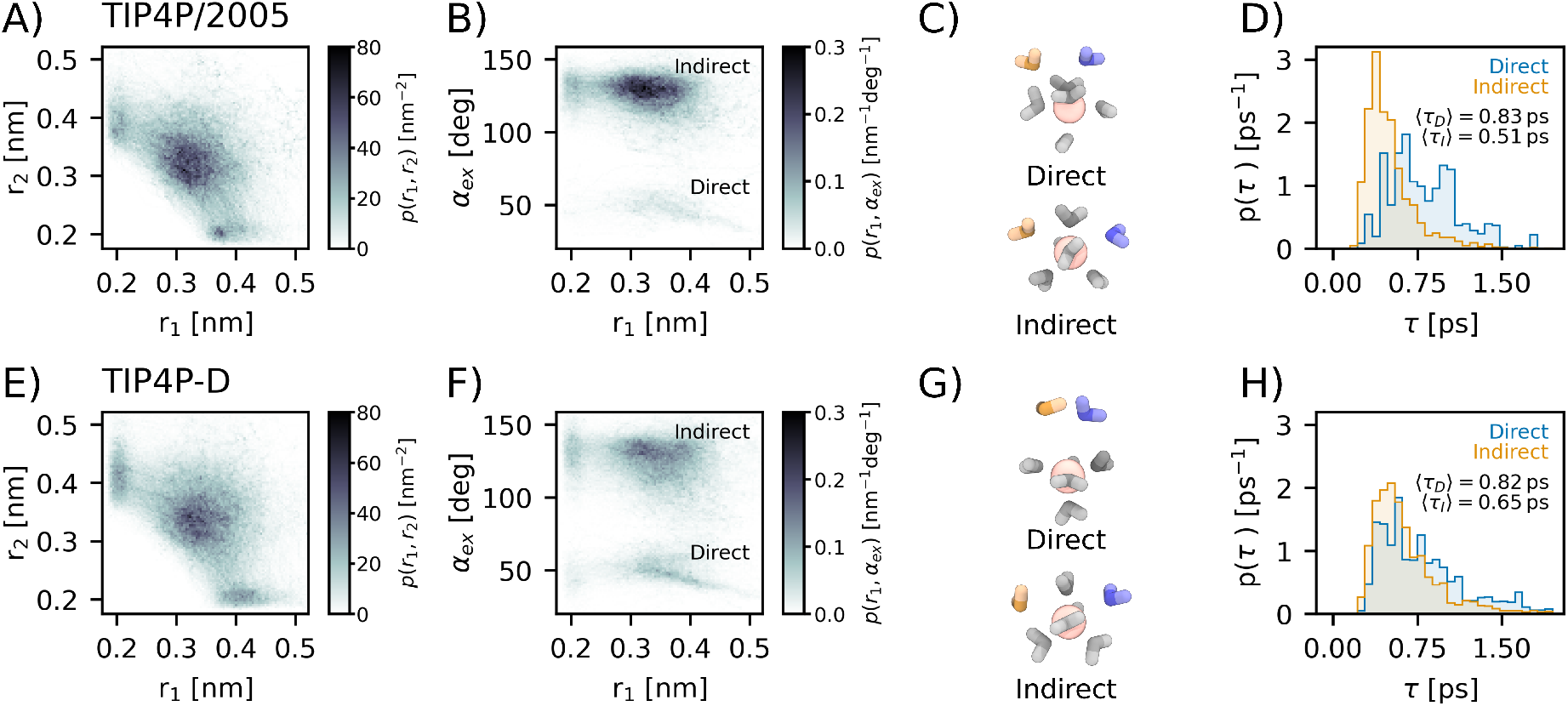
Water exchange with the four-site water models TIP4P/2005 (top) and TIP4P-D (bottom). (A,E) Probability distribution function *p*(*r*_1_, *r*_2_) along transition pathways in dependence of the distances *r*_1_ and *r*_2_. (B, F) Probability distribution function *p*(*r*_1_, *α*_ex_) in dependence of *r*_1_ and the exchange angle *α*_ex_. (C,G) Representative simulation snapshots of the transition state for the direct and indirect mechanism. (D, H) Distribution of transition times *p*(*τ*) for direct and indirect exchange pathways.

**FIG. 4:**
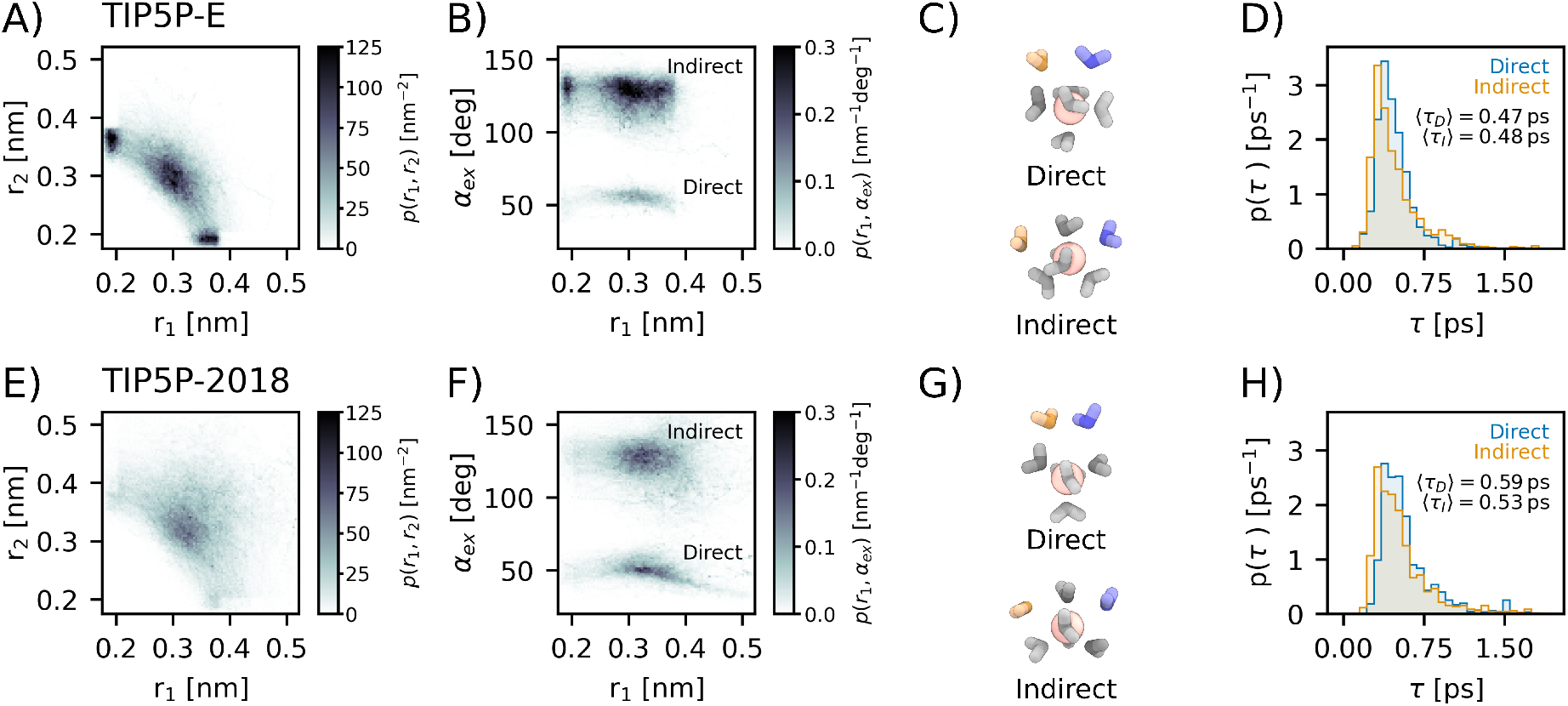
Water exchange with the five-site water models TIP5P-E (top) and TIP5P/2018 (bottom). (A,E) Probability distribution function *p*(*r*_1_, *r*_2_) along transition pathways in dependence of the distances *r*_1_ and *r*_2_. (B, F) Probability distribution function *p*(*r*_1_, *α*_ex_) in dependence of *r*_1_ and the exchange angle *α*_ex_. (C,G) Representative simulation snapshots of the transition state for the direct and indirect mechanism. (D, H) Distribution of transition times *p*(*τ*) for direct and indirect exchange pathways.

### Interchange associative pathways for polarizable and long-ranged force fields

Interestingly, an interchange associative exchange mechanism is observed with the polarizable AMOEBA force field and with the recently developed non-polarizable microMg parameters. The distributions of exchange distances along reactive pathways (Figure 5A,B and Figure 5E,F) show that the exchange takes place inside the first hydration shell (Figure S1) and the distances in the transition state are only slightly larger compared to their equilibrium values (Table I). Similar to the dissociative pathways, direct and indirect exchanges are observed. Here, water molecules exchange in an angle of approximately 70° or 137° for direct and indirect paths, respectively. With seven water molecules in the inner shell, the transition state resembles a pentagonal bipyramidal geometry (Figure 5C, G). Consequently, the exchange angle is approximately a multiple of the central 72° angle. Considering the permutation invariance of solvent molecules, the transition states of direct and indirect reactions become virtually indistinguishable.

**FIG. 5:**
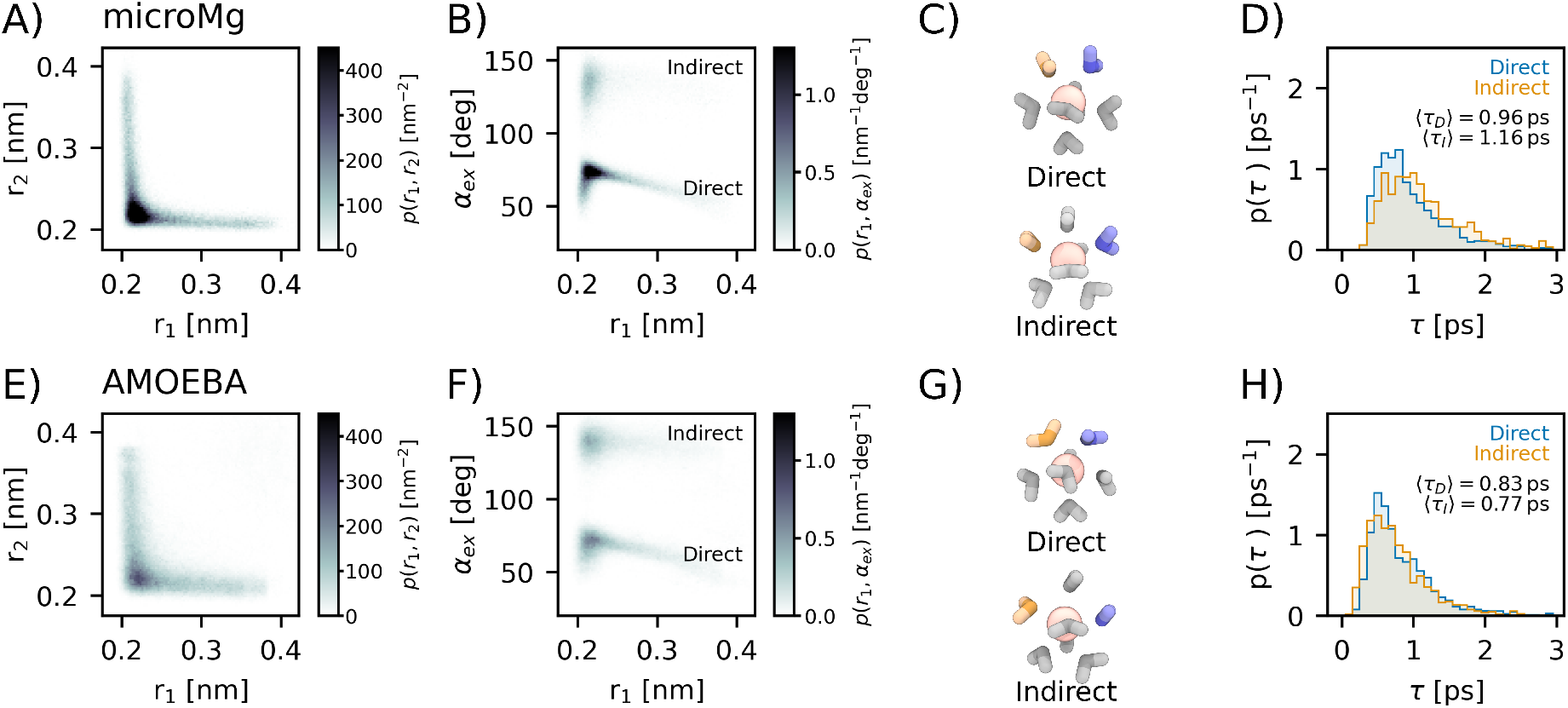
Water exchange with the microMg parameters and TIP3P (top) and with the polarizable AMOEBA force field (bottom). (A,E) Probability distribution function *p*(*r*_1_, *r*_2_) along transition pathways in dependence of the distances *r*_1_ and *r*_2_. (B, F) Probability distribution function *p*(*r*_1_, *α*_ex_) in dependence of *r*_1_ and the exchange angle *α*_ex_. (C,G) Representative simulation snapshots of the transition state for the direct and indirect mechanism. (D, H) Distribution of transition times *p*(*τ*) for direct and indirect exchange pathways.

The reason for the crossover from dissociative to associative mechanism is the range of the atomistic Mg^2+^-water interaction potential of the different force fields. Due to their high charge density, Mg^2+^ ions polarize their environment strongly. Consequently, charge-induced dipole interactions and charge transfer can become significant and render the Mg^2+^-water interactions more attractive and long-ranged. The AMOEBA force field takes some of these many-body quantum effects into account explicitly while the microMg parameters account for them implicitly. In both cases, the Mg^2+^-water interaction potentials are more long-ranged compared to standard force field parameters and have a non-vanishing attractive contribution at the edge of the first hydration shell (Figure S2). Consequently, in the transition state, the exchanging waters have considerable interactions with the Mg^2+^ ion leading to the observed I_*a*_ mechanism. By contrast, for the more short-ranged interaction potential in the Mamatkulov Mg^2+^ force field, the interactions of the entering and leaving waters with Mg^2+^ are almost negligible.

The crossover from interchange dissociative to associative also affects the exchange rate. For the standard Mg^2+^ force field in combination with TIP3P water, the rate constant is *k* = 24.0 *±* 8.8 s^−1^ [12] and significantly lower than the experimental result (*k* = 5.3 *·* 10^5^ s^−1^ [6]). With the microMg parameters, the rate constant is *k* = 8.0 *±* 1.2 *·* 10^5^ s^−1^ and closely matches the experimental value [13]. The polarizable AMOEBA force field yields the highest rate *k* = 0.5 *·* 10^9^ s^−1^ [10, 27] and significantly overestimates the experimental result.

## CONCLUSION

Water exchange between the hydration shells of Mg^2+^ is essential for a large variety of physiological processes. In order to correctly capture the exchange dynamics in all-atom molecular dynamics simulations, accurate force fields are required. To ease the choice of the force field, we have systematically investigated the influence of different water models and Mg^2+^ force fields on the mechanism of water exchange. In particular, we have used transition path sampling as a particularly powerful sampling strategy to provide unbiased insights into the kinetic pathways and to cover the long time scales involved in water exchange.

In all cases, water exchange can occur either via an indirect or direct exchange pathway (exchanging molecules occupy different/same position on the water octahedron) without stable intermediates. This provides further evidence that the I_*d*_ mechanism is dominant while dissociative (D) pathways with a stable, reduced coordination intermediates can be excluded, in agreement with quantum mechanical calculations in the gas-phase [42]. In addition, a crossover from an interchange dissociative (I_*d*_) to an associative (I_*a*_) reaction mechanism is observed dependent on the range of the Mg^2+^-water interaction potential of the respective force field.

No force field combination tested in our current work is able to simultaneously reproduce the experimentally observed I_*d*_ mechanism and the rate of water exchange. Standard nonpolarizable Mg^2+^ force fields in combination with the commonly used rigid water models yield the I_*d*_ mechanism in agreement with experimental results but significantly underestimate the exchange rate [12, 13, 19, 20]. The microMg parameters [13] yield close agreement with the experimental rate but follow the I_*a*_ mechanism in contrast to the experimental results. Finally, the polarizable AMOEBA force field [28, 29] significantly overestimates the experimental rate and yields the I_*a*_ mechanism.

Consequently, the microMg force field parameters [13] should be used in bimolecular simulations addressing the binding kinetics and the Mamatkulov Mg^2+^ parameters [20] when addressing the mechanism of ligand exchange reactions. The applicability of the polarizable AMOEBA force field is clearly limited. However, models with variable polarizability [10] depending on the distance between Mg^2+^ and water might improve the agreement with experimental results.

In any case, the impact of water model and ionic force field must be explored in more detail in order to develop a more accurate description. In particular, designing an experiment that could probe water exchange pathways directly would be an invaluable contribution to the field. Alternatively, ab initio QM/MM molecular dynamics simulations [11] at high levels of accuracy could provide further insights and a step toward improved models for biomolecular simulations.

## SUPPORTING INFORMATION

The data that support the findings of this study are available from the corresponding author upon reasonable request. See the supplementary material for further discussion of the radial distribution function and the Mg^2+^-water interaction potentials.

## ACKNOWLEDGMENTS

This work was funded by the Deutsche Forschungsgemeinschaft (DFG, German Research Foundation), Emmy Noether program, 315221747. LOEWE CSC and GOETHE HLR are acknowledged for supercomputing access.

## SUPPORTING INFORMATION

### RADIAL DISTRIBUTION FUNCTION

The radial distribution functions were calculated from MD simulations in the NPT ensemble. For non-polarizable force fields, simulations were carried out up to a length of 10 ns. Afterwards, the radial distribution functions were obtained using Gromacs 2018.8 [1] with a binning of 0.002 nm. For the polarizable AMOEBA force field, a 10 ns simulation was performed and the radial distribution function was calculated using MDAnalysis [2, 3] with a binning of 0.002 nm.

**FIG. S1.**
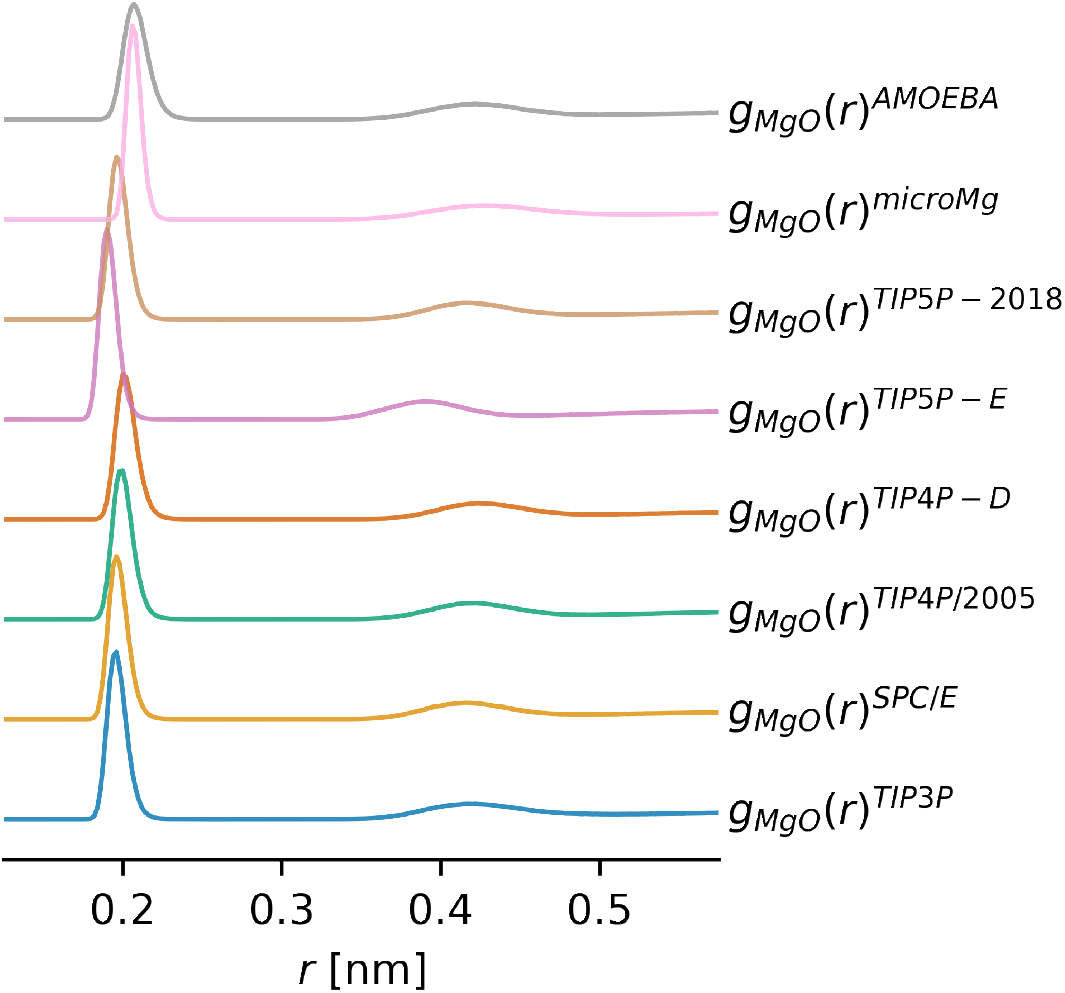
Radial distribution functions g_MgO_(r) for different water models and Mg^2+^ force fields.

### INTERACTION POTENTIALS

Interaction potentials between Mg^2+^ and water were calculated in a simplified system. For all water models, Mg^2+^ and the water’s oxygen and hydrogen atoms were placed in the x/y plane. The rotation of the water molecule was fixed so that both hydrogen atoms were equidistant from the Mg^2+^ ion. In the case of four- and five-site water models, the massless charge sites and lone pairs were placed relative to the oxygen and hydrogen atoms. For non-polarizable force fields, the interaction energy was evaluated for different Mg^2+^-OW distances in steps of 0.005 nm using Gromacs 2018.8. For the AMOEBA force field, the interaction energy was estimated using OpenMM 7.4.1 [4]. Here, the potential energy was first calculated for a system with Mg^2+^ and water at a certain distance. Subsequently, the potential energy of a system containing only Mg^2+^ and only water was subtracted to obtain the interaction potential.

**FIG. S2.**
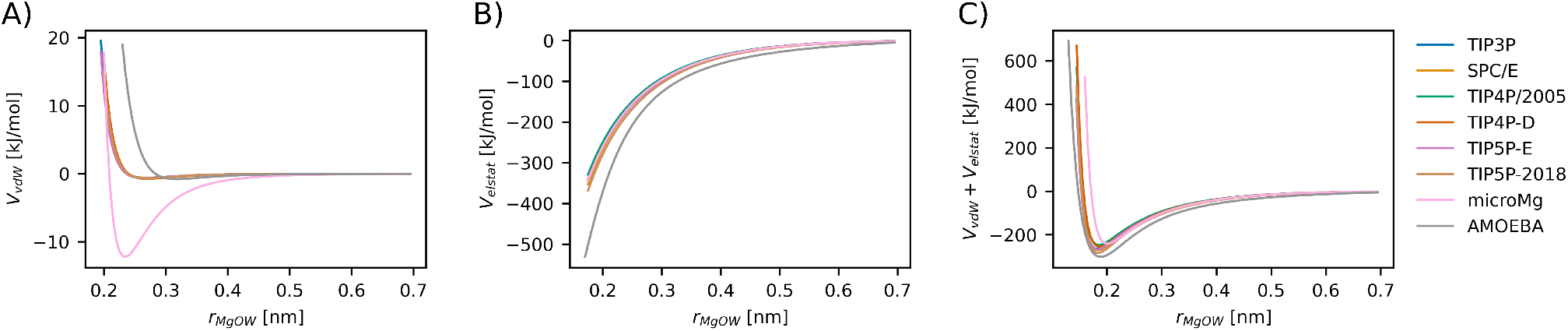
Interaction potentials between water and Mg^2+^ as a function of the Mg^2+^-OW distance for different water models and Mg^2+^ force fields. The interaction potential is decomposed into (A) Van der Waals and (B) electrostatic contributions. The combined interaction potential is shown in (C).

## Notes

### Competing Interest Statement

The authors have declared no competing interest.

